# Cord blood buffy coat DNA methylation is comparable to whole cord blood methylation

**DOI:** 10.1101/181057

**Authors:** John Dou, Rebecca J. Schmidt, Kelly S. Benke, Craig Newschaffer, Irva Hertz-Picciotto, Lisa A. Croen, Ana-Maria Iosif, Janine M. LaSalle, M. Daniele Fallin, Kelly M. Bakulski

**Author notes:** **Disclosures of interest**: The authors report no conflict of interest.

## Abstract

**Background:** Cord blood DNA methylation is associated with numerous health outcomes and environmental exposures. Whole cord blood DNA reflects all nucleated blood cell types, while centrifuging whole blood separates red blood cells by generating a white blood cell buffy coat. Both sample types are used in DNA methylation studies. Cell types have unique methylation patterns and processing can impact cell distributions, which may influence comparability.

**Objectives:** To evaluate differences in cell composition and DNA methylation between buffy coat and whole cord blood samples.

**Methods:** Cord blood DNA methylation was measured with the Infinium EPIC BeadChip (Illumina) in 8 individuals, each contributing buffy coat and whole blood samples. We analyzed principal components (PC) of methylation, performed hierarchical clustering, and computed correlations of mean-centered methylation between pairs. We conducted moderated t-tests on single sites and estimated cell composition.

**Results:** DNA methylation PCs were associated with individual (*P*_PC1_=1.4x10^-9^; *P*_PC2_=2.9x10^-5^; *P*_PC3_=3.8x10^-5^; *P*_PC4_=4.2x10^-6^; *P*_PC5_=9.9x10^-13^), and not with sample type (P_PC1-5_>0.7). Samples hierarchically clustered by individual. Pearson correlations of mean-centered methylation between paired individual samples ranged from r=0.66 to r=0.87. No individual site significantly differed between buffy coat and whole cord blood when adjusting for multiple comparisons (5 sites had unadjusted *P*<10^-5^). Estimated cell type proportions did not differ by sample type (P=0.86), and estimated cell counts were highly correlated between paired samples (r=0.99).

**Conclusions:** Differences in methylation and cell composition between buffy coat and whole cord blood are much lower than inter-individual variation, demonstrating that both sample preparation types can be analytically combined and compared.

## Introduction

Early life epigenetic epidemiology is a highly promising and productive area of research ^1-3^. Prenatal environmental exposures may influence epigenetic factors such as DNA methylation, which serves as a useful biomarker of previous exposures and conditions ^4,5^. Similarly, DNA methylation at birth is also associated with birth outcomes and may mediate impacts on health outcomes later in life ^6^. Biological samples collected at birth allow for the investigation of epidemiological links between exposures, epigenetic changes through DNA methylation, and health. Cord blood is often used as a proxy tissue in methylation studies on infants. Cord blood DNA methylation has been associated with numerous health outcomes and environmental exposures ^7-13^.

Cord blood contains a mixture of DNA from multiple leukocyte cell types including granulocytes, other white blood cells, and nucleated red blood cells present in fetal life. In contrast, non-nucleated red blood cells in blood do not contribute DNA to methylation measures. Since cord blood cell types have unique methylation patterns ^14-16^, differential cell proportions across samples highly influence DNA methylation measures and can confound analyses if related to variables of interest ^17^.

Critical to epidemiological testing and interpretation, blood sample processing impacts cell composition. DNA isolations from whole cord blood, commonly collected in anti-coagulant tubes (containing heparin or EDTA), reflect all nucleated cell types, including nucleated red blood cells. Some investigators elect to centrifuge the whole blood, generating a buffy coat of white blood cells (including granulocytes) separated from red blood cells. These two methods are the most commonly used for processing cord blood in epigenetic epidemiology. A third processing method, beyond the scope of this study, involves density centrifugation (e.g. with Ficoll), to isolate cord blood mononuclear cells (CBMCs), removing both granulocytes and red blood cells ^18^. Uniquely hypomethylated fetal nucleated red blood cells have adhesive properties^15^. We hypothesize that buffy coat separation may not successfully remove this cell type.

Replication across studies is an essential component of epigenetic epidemiology, but the comparability across sample processing methods has not yet been tested. Therefore, it is important to experimentally test the effect of cord blood sample processing on the cellular composition and thus the DNA methylation of samples commonly used for epidemiological measures. We examined differences in cell proportions and DNA methylation by whole cord blood or cord buffy coat sample type. We hypothesized that since nucleated red blood cells stick to white blood cells, buffy coat samples will be similar in cell composition and DNA methylation to whole cord blood samples.

## Methods

### Study sample

The Early Autism Risk Longitudinal Investigation (EARLI) is a pregnancy cohort study that recruited mothers who already had a child diagnosed with an autism spectrum disorder. Full details of the study have been previously described ^19^. Informed consent was obtained from all participants. This study was approved by Institutional Review Boards at all study sites (Johns Hopkins University, Drexel University, UC Davis, Kaiser Permanente) and secondary analysis site (University of Michigan). Demographic characteristics were self-reported throughout pregnancy. At birth, biological samples including cord blood were collected. In EARLI, whole cord blood was drawn into EDTA tubes. One tube was archived as whole blood and the second was centrifuged to separate the buffy coat. Samples were archived at -80°C prior to processing for this study. For this sub-study, we randomly selected eight term births with adequate buffy coat and whole blood biorepository samples.

### DNA methylation measures

DNA was extracted using the DNeasy Blood kit (Qiagen) per manufacturer instructions. DNA was bisulfite treated and cleaned using the EZ DNA methylation kit (Zymo Research) at the University of Michigan Epigenetics Core. Bisulfite treated DNA was hybridized and imaged on the Infinium MethylationEPIC (EPIC) BeadChip (Illumina) at the University of Michigan DNA Sequencing Core. Samples were randomly plated to reduce potential batch effects. In total, there were 8 whole cord blood DNA paired with 8 buffy DNA from the same cord sample among EARLI participants. Raw DNA methylation data are available through the Genome Expression Omnibus (GEO pending).

### Data preprocessing

Raw image array files were processed in R statistical software (version 3.3) using the minfi package (version 1.20.2) ^20^. We estimated sex from raw methylation data, and observed pair concordance with demographic data. Probes on sex chromosomes were included in the analysis. All samples had overall methylated and unmethylated image intensities greater than 11 relative fluorescence units, and fewer than 1% failed probes. Data were processed with noob background correction ^21^. Probes that failed detection p > 0.01 in 1 or more samples were dropped (n = 2,699). An additional 43,096 probes annotated as cross reactive were also removed ^22^. The final study sample contained 821,041 probes from 16 samples. Beta values from the preprocessed data were used to approximate percent methylation per site.

### Statistical analyses

We examined DNA methylation distributions of samples using density plots of beta values, colored by sample pair and sample type. DNA methylation data were compared using principal components analysis (PCA). In pairwise PCA plots up to the sixth principal component (PC), data points were colored by technical covariates (slide, row number), biological covariates (sex, race), and experimental conditions (individual, sample type) to visualize differences by these variables. We tested for these differences in PCA using t-tests for categorical variables, ANOVA for multi-category data, and Pearson correlation tests for continuous variables. We then performed hierarchical clustering on the DNA methylation data, with Euclidian distance calculation and complete linkage clustering. Cord blood cell composition was estimated from DNA methylation ^14^. We compared estimated cell distributions by sample type (buffy coat, whole blood) using ANOVA and Pearson correlation.

DNA methylation in whole blood and buffy coat were compared using plots of mean-centered correlations at all sites for each pair ^23^. To center data for each CpG, the mean across all samples was subtracted from the observed values in each sample for that CpG. Pearson correlations were then computed between paired buffy coat and whole blood samples ^23^. We also created Bland and Altman plots ^24^ to evaluate differences between methylation values measured from paired samples.

Single site analysis was performed to investigate possible CpGs that were differentially methylated in buffy coat and whole blood samples. We performed moderated paired t-tests for each CpG using the limma package (version 3.30.13) ^25^. No other covariates were included in the model for adjustment, due to the paired design. When adjusting for multiple comparisons, we considered *P* < 6.09 x 10^-8^ reaching genome-wide significance. We compared these results to published signatures of cord blood cell types ^14^.

## Results

### Study sample characteristics

Among the 8 participants in our study (16 paired whole blood and buffy coat samples), 3 infants were female, 6 mothers identified as white, 1 as Asian Indian, and 1 as mixed race (**Table 1**). Mean gestational age at birth was 38.9 weeks (standard deviation=0.6).

**Table 1.** Subject and Sample Characteristics

| **Table 1. Subject and Sample Characteristics** |  |  |  |
| --- | --- | --- | --- |
| Sex (N, %) |  |  |  |
| Male |  | 5 (62.5%) |  |
| Female |  | 3 (37.5%) |  |
| Race (N, %)<sup>a</sup> |  |  |  |
| White |  | 6 (75%) |  |
| Asian Indian |  | 1 (12.5%) |  |
| More than one |  | 1 (12.5%) |  |
| Gestational Age (mean, IQR) |  |  |  |
| Weeks |  | 38.86 (0.55) |  |
| Cell Proportions (mean, IQR) | Buffy Coat | Whole Blood | p-value<sup>c</sup> |
| CD8T | 11.3% (2.8%) | 11.7% (4.6%) | 0.86 |
| CD4T | 17.3% (4.2%) | 16.0% (4.2%) |  |
| NK | 0.2% (0%)<sup>b</sup> | 0.3% (0%)<sup>b</sup> |  |
| Bcell | 9.2% (2.3%) | 9.5% (2.8%) |  |
| Mono | 8.9% (2.7%) | 9.4% (3.6%) |  |
| Gran | 47.3% (16.5%) | 47.7% (10.1%) |  |
| nRBC | 7.4% (4.5%) | 8.3% (5.5%) |  |

<sup>a</sup> Self-reported from questionnaire
<sup>b</sup> Majority of samples had predicted natural killer count of 0
<sup>c</sup> ANOVA for proportions by sample type and cell type, p-value for sample type shown

### Principal components of DNA methylation are associated with individual, not sample type

Overall distributions of DNA methylation by sample did not group by buffy coat or whole blood sample type (**Supplementary Fig 1**). In PCA, samples separated by individual rather than sample type (buffy coat/whole blood), suggesting there were greater differences between individuals than within (**Figure 1**). We observed differences by the technical variable slide (**Supplementary Figure 2**), as well as differences by sex and race (**Supplementary Figure 3**).In hierarchical clustering of DNA methylation data, samples also grouped by individual rather than type (**Figure 2**). Principal component (PC) 1 of the DNA methylation data was associated with individual (*P* = 1.4x10^-9^), sex (*P* = 4.2x10^-9^), estimated percent granulocytes (*P* = 1.08x10^-4^), and estimated percent nucleated red blood cells (*P* = 6.0x10^-5^) (**Supplementary Figure 4**).PC 2 was associated with individual (*P* = 2.9x10^-5^) and slide (*P* = 5.3x10^-3^). PCs 3-5 were further associated with individual (*P* = 3.8x10^-5^, 4.2x10^-6^, 9.9x10^-13^, respectively). Importantly, sample type (buffy versus whole) was not associated with any DNA methylation PC, with all *P* > 0.7 (**Supplementary Figure 4**).

**Figure 1.**
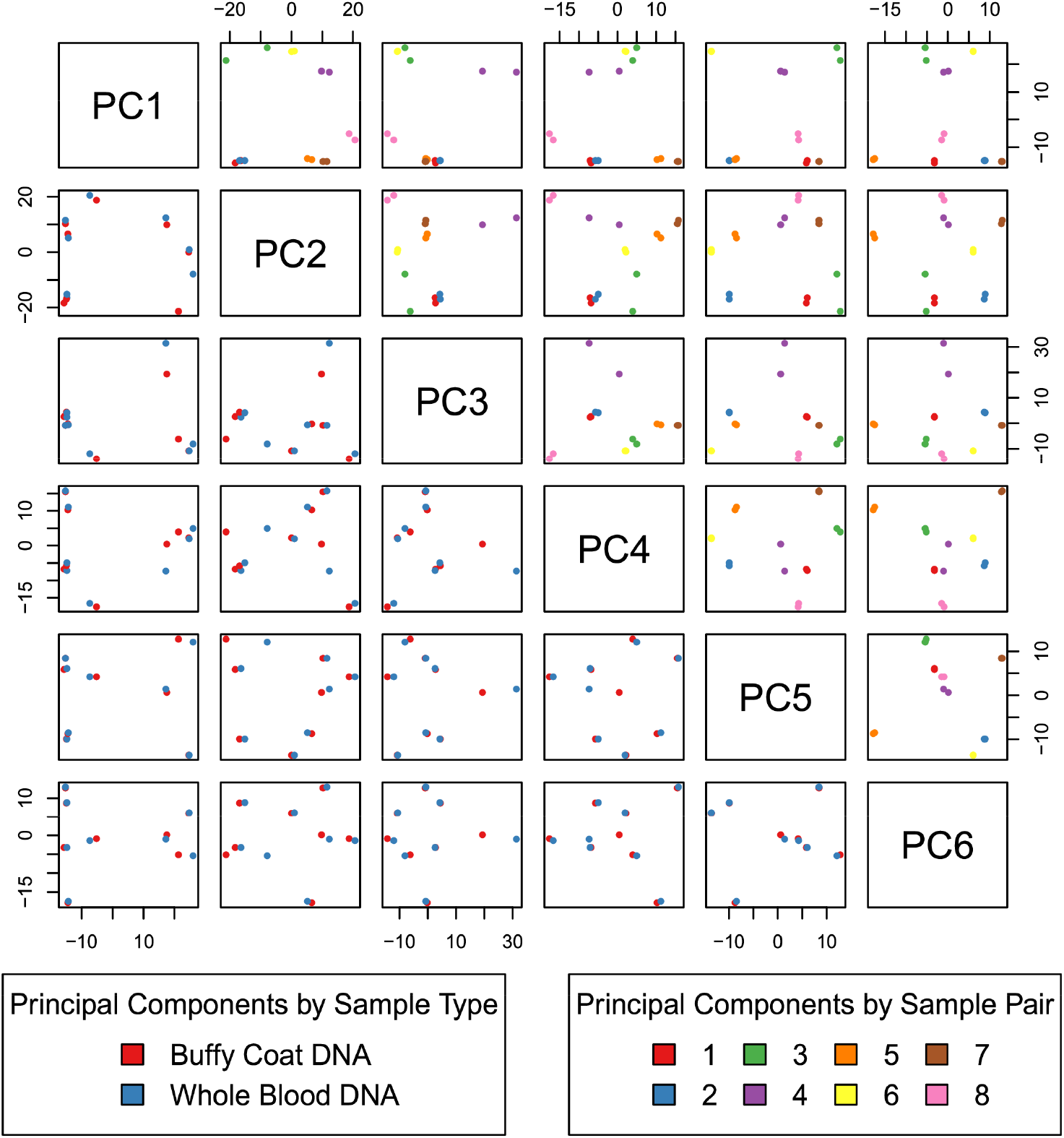
Principal components analysis of EPIC array DNA methylation for 16 cord blood samples (pairs of 8 buffy coat and 8 whole blood). We do not observe differences by sample type (blue: buffy coat, red: whole blood) and paired samples tended to cluster together.

**Figure 2.**
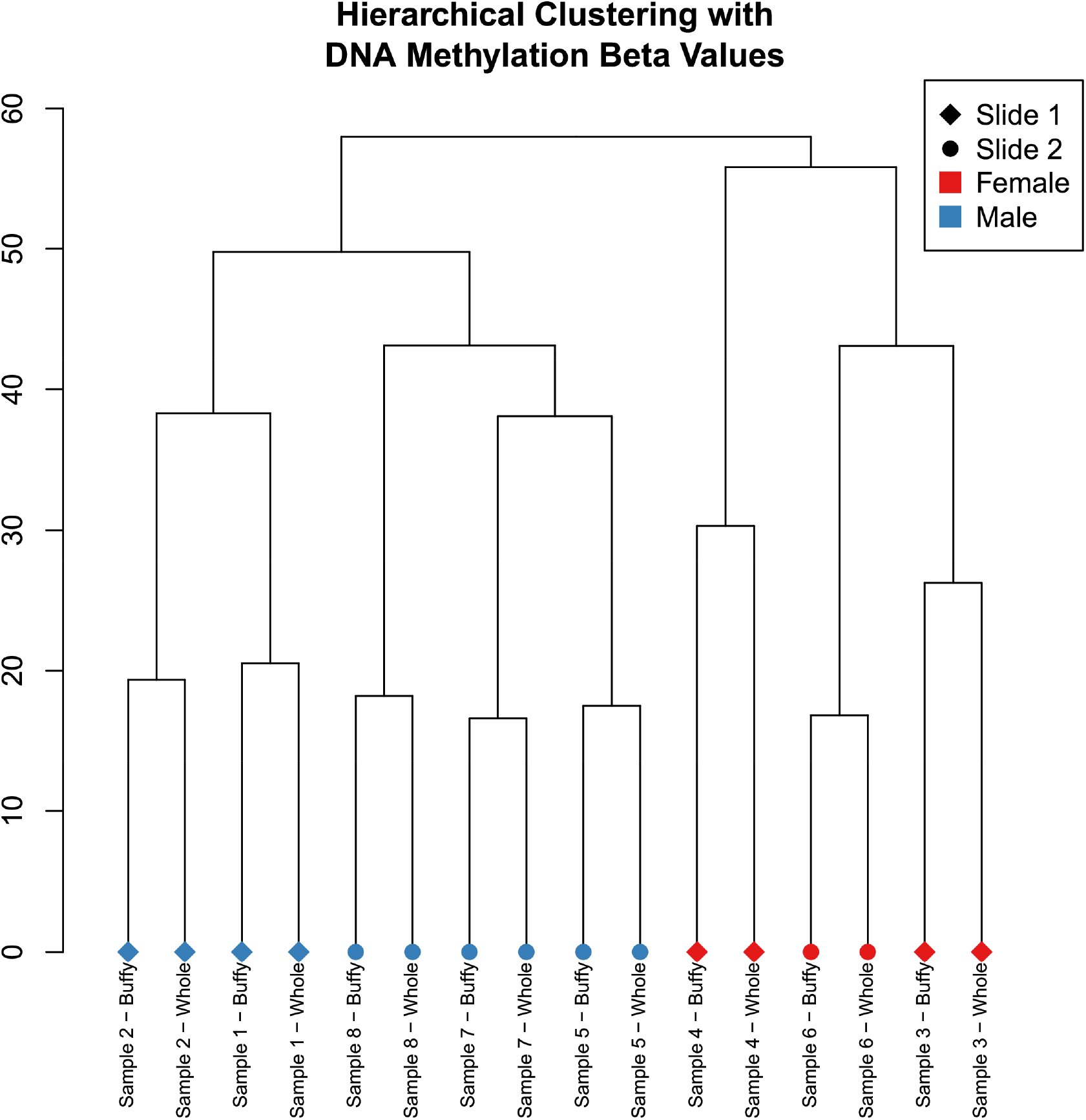
Dendrogram from hierarchical clustering on DNA methylation data, using complete linkage method. Samples clustered together by pairs.

### Whole cord blood and buffy coat DNA methylation measures are correlated within individuals

Individual pairs of whole blood and buffy coat mean-centered DNA methylation values were highly correlated (**Figure 3**). The least correlated sample pair had Pearson r = 0.66, while the most correlated had r = 0.87. As seen in Bland-Altman plots, the differences in methylation values tended to be small and were scattered around 0, with a handful of exceptions in each pair with extreme differences (**Supplementary Figure 5**). When characterizing variation between the pairs, 50% of absolute differences were within approximately 1% methylation, and 95% of absolute differences were within approximately 5% methylation.

**Figure 3.**
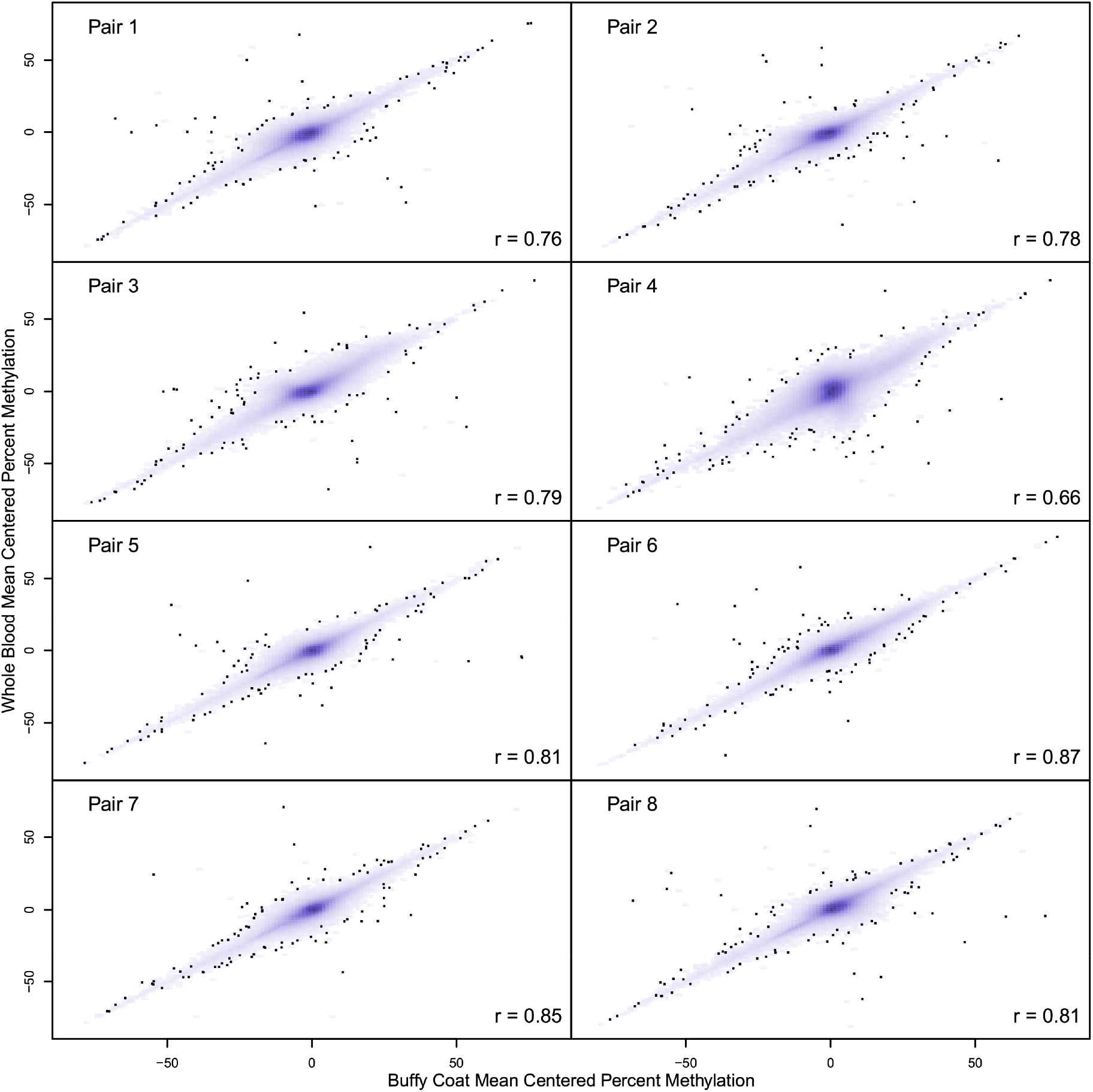
Mean centered percent methylations at CpGs in buffy coat and whole blood paired samples. 100 points in the least dense regions are plotted. We observe high correlation between methylation in buffy coat and whole blood.

### Estimated cell proportions do not differ between buffy coat and whole cord blood

Cell proportions were estimated for each sample (**Figure 4**). Granulocytes made up the largest fraction, with estimated proportions ranging from 35% to 60%. Natural killer cells had the smallest fractions, with most samples (12 of 16) having an estimated proportion of 0%. Buffy coat did indeed include nucleated red blood cells in cell type estimation (mean = 7.4%). We did not observe differences in estimated cell proportions between buffy coat and whole blood samples (*P* = 0.86) (**Table 1**). Estimated cell proportions were highly correlated (Pearson r = 995). Within individual cell types, correlation was very high between buffy coat and whole cord blood samples (CD8T r = 0.97, CD4T r = 0.97, B cell r = 0.93, monocytes r = 0.94, granulocytes r = 0.96, nRBC r = 0.99, NK r = 1 due to previously mentioned 0% estimates).

**Figure 4.**
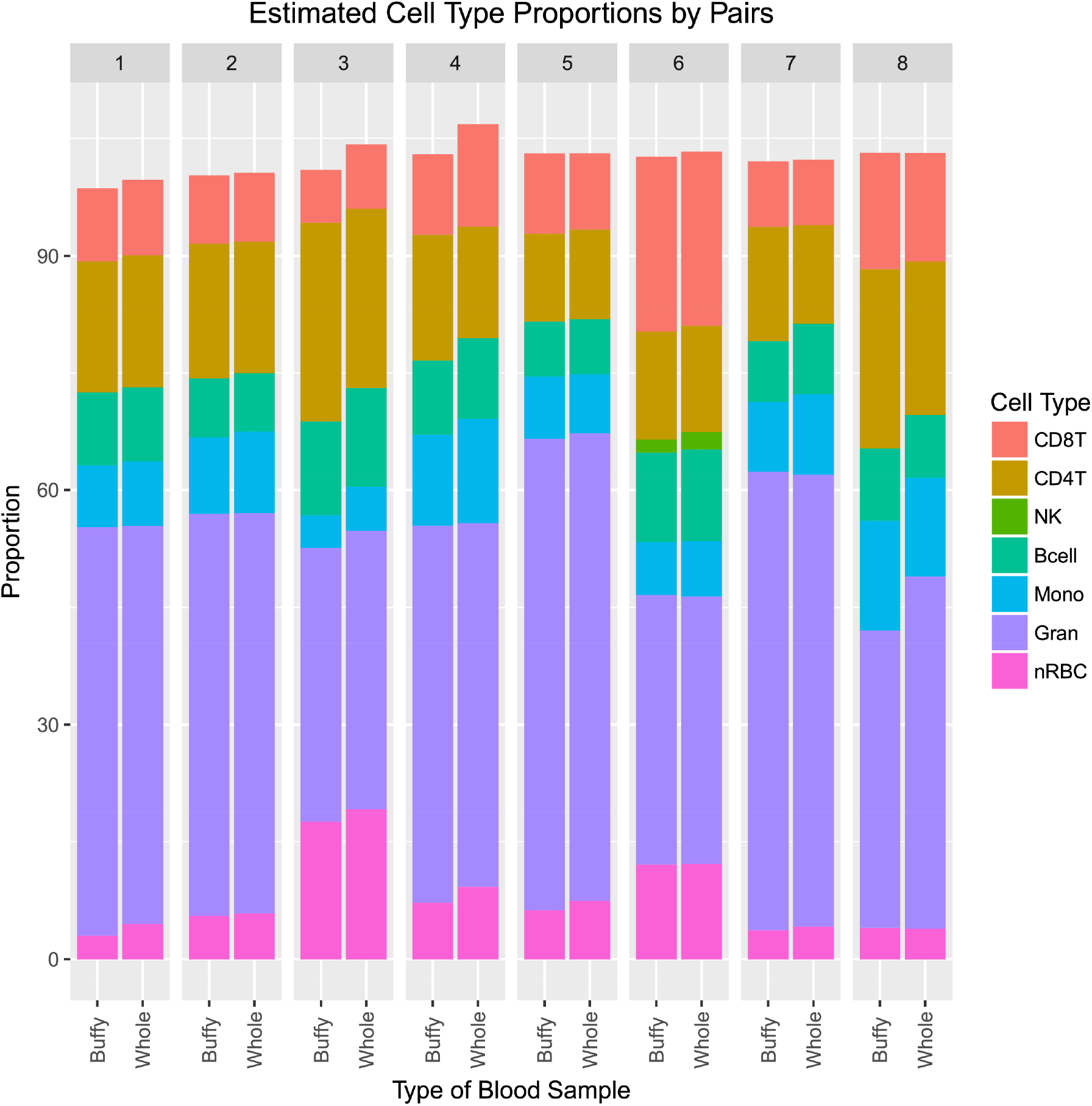
Estimated cell type proportions from 8 matched buffy coat and whole blood cord samples. We observe that cell type proportions within subject are correlated.

### Individual CpG sites are not associated with buffy versus whole cord blood sample type

In single site analysis, no CpG site was significant when adjusting for multiple comparisons. The model had a lambda inflation factor of 1.37 (**Supplementary Figure 6**). The lowest observed unadjusted P-value was 1.63 x 10^-6^ for cg10480693, which had an estimated mean difference of 2.69% methylation comparing whole cord blood to buffy coat. Five sites had *P* < 10^-5^ (summarized in **Supplementary Table 1**, results for all probes can be found in **Supplementary Table 2**), adjusting for false discovery rate yielded Q-values of 0.66 for these sites. Three of these five sites were not on 450k arrays used in previous literature on individual cord blood sorted cell types. Among those overlapping with the 450k array, cg00648883 was previously strongly associated with cell type and was hypomethylated in monocytes and granulocytes relative to other cell types ^14^. Further, cg10701033 was slightly hypomethylated in nucleated red blood cells ^14^.

## Discussion

Paired samples of whole cord blood and buffy coat had similar DNA methylation estimated cell type proportions, including nucleated red blood cells. Methylation measures were correlated between paired buffy coat and whole blood samples, with Pearson correlation coefficients ranging from 0.66 to 0.87. Principal components of DNA methylation show that the samples cluster by subject, and not by sample type. These findings have important implications for whole cord blood and buffy coat sample DNA methylation comparability. We observed that DNA methylation and cell composition differences between whole blood and buffy coat sample types are minimal, and we therefore predict that epigenome-wide association studies may replicate across these sample types.

Newborn cord blood has often been used in epigenetic epidemiology studies. In whole cord blood, DNA methylation associations were identified with prenatal antidepressant exposure ^26^, fetal growth restriction ^27^, and cord blood triglyceride levels ^28^. Other studies elected to process and extract DNA from buffy coat, and identified associations between DNA methylation and maternal depression or anxiety ^29^, parental obesity ^30^, birth weight-for-gestational age ^13^, and prenatal mercury exposure ^31^. Whole cord blood and cord blood buffy coat are the most commonly used sample types in epigenetic epidemiology.

Meta-analyses and replication testing is an important and necessary future direction for epigenetic epidemiology to overcome sample size limitations with multiple comparisons. In general, we should take care when combining findings from samples with different underlying cell distributions. Differences in nucleated red blood cell counts are especially worrisome, as they have been observed to have some of the most extensive methylation changes among the cell types in cord blood ^12^. Specific to cord blood buffy coat and whole blood samples, our findings of similar cell compositions between the two sample types gives some reassurance to this concern. Indeed, studies have meta-analyzed epigenetics data from both whole cord blood and buffy coat sources. For example, in a meta-analysis of maternal smoking and newborn cord blood DNA methylation from 13 cohorts, 2 used buffy coat samples and 11 whole blood ^7^.

Observed associations between CpG methylation and maternal smoking were generally consistent across cohorts, regardless of sample type ^7^. Replication across studies is an essential component of genome-wide testing and whole cord blood and buffy coat are compatible sample types.

One limitation to our study was the relatively small sample size with only 8 buffy coat and whole cord blood pairs, which limits power to detect small differences that may exist. The pairing of samples types by individual subject alleviates some of this weakness. Pairing also controls for other factors that impact DNA methylation, such as gestational age. Our samples were limited to full term cord blood, and preterm infants display altered cord blood DNA methylation, including differences in cell type composition ^12^. Future work should compare DNA methylation of blood sample types in preterm infants. The Illumina EPIC array is a widely used platform, and its use in this study offers coverage across the genome at approximately 850,000 sites. DNA was isolated in all samples using the DNeasy Blood kit (Qiagen), but use of different methods for isolation may impact DNA methylation. Another potential limitation of this study is that we did not evaluate other blood sample processing methods, such as using Ficoll density centrifugation to isolate cord blood mononuclear cells (CBMCs) ^18^. Since Ficoll separates out granulocytes, it is likely these sample types would have different cell type distributions and methylation measurements. Further testing should investigate PBMCs relative to whole cord blood and buffy coat.

Our results suggest that DNA methylation measurements from buffy coat and whole cord blood samples are highly comparable on the new EPIC array platform. DNA methylation was highly correlated between the two sample types, and principal components of methylation data reveal clustering by sample pairs. Differences between whole cord blood and buffy coat are much smaller than the inter-individual differences, and the frequency of nucleated red blood cells is not significantly different by sample processing. Thus future studies, and those that have already done so, can accurately combine and compare results from buffy coat and whole cord blood samples.

## Acknowledgments

We thank the EARLI study participants and staff. We thank the Johns Hopkins Biological Repository (JHBR) for archiving and processing the samples. We thank the University of Michigan Epigenetics and DNA Sequencing Cores for conducting the DNA methylation measurements.

